# Design of a Toolbox of RNA Thermometers

**DOI:** 10.1101/017269

**Authors:** Shaunak Sen, Divyansh Apurva, Rohit Satija, Dan Siegal, Richard M Murray

## Abstract

Biomolecular temperature sensors can be used for efficient control of large-volume bioreactors, for spatiotemporal control and imaging of gene expression, as well as to engineer robustness to temperature in biomolecular circuit design. While RNA-based sensors, called ‘thermometers’, have been investigated in natural and synthetic contexts, an important challenge is to design different responses to temperature, differing in sensitivities and thresholds. We address this issue using experimental measurements in cells and in cell-free biomolecular ‘breadboards’ in combination with computations of RNA thermodynamics. We designed a library of RNA thermometers, finding, computationally, that it could contain a multiplicity of responses to temperature. We constructed this library and found a wide range of responses to temperature, ranging from 3.5-fold to over 10-fold in the temperature range 29°C – 37°C. These were largely linear responses with over 10-fold difference in slopes. We correlated the measured responses with computational expectations, finding that while there was no strong correlation in the individual values, the overall trends were similar. These results present a toolbox of RNA-based circuit elements with varying temperature sensitivities.

## 1 Introduction

Biomolecular temperature sensors, which convert temperature into a biologically functional response, can have multiple applications. These include, for example, large-volume bioreactors, where such sensors allow the use of heat as an inducer, which may be more convenient than a chemical-based inducer. As another example, these sensors can be used to spatiotemporally control pathways inside cells and tissues through local heating induced by electromagnetic waves in the millimetre range [12]. In the case that these regulate the expression of reporters, they can be used for spatiotemporally precise imaging as well. Finally, temperature sensors may find application in engineering temperature compensation in biomolecular circuit design, where it is often desired to have specifically-tailored responses to temperature to counteract, and hence cancel, other changes with temperature [5,11]. To meet these needs, it is desirable to have a toolbox of sensors with different sensitivities and thresholds in their respective temperature responses.

Temperature-sensing RNA molecules, called ‘RNA thermometers’, operate in diverse natural contexts, mediating cellular behaviours such as heat-shock response and pathogenesis [6]. There are many mechanisms for the operation of an RNA thermometer, each of which depend on a temperature-dependent conformational change. At its simplest, a RNA thermometer consists of an RNA sequence containing the ribosome binding site (RBS, Fig. 1A). At a permissive temperature, the ribosome can access this RBS and translation proceeds efficiently. At a restrictive temperature, the RNA sequence folds in such a manner so as to prevent this access and inhibit translation. Examples include the *rpoH* gene mediating heat-shock response in *E. coli* and the *cIII* gene regulating the life cycle of phage λ. While these natural examples have a relatively complicated secondary structure with multiple stems, hairpin loops, and bulges, RNA thermometers have been designed with simpler secondary structures, with only a single stem-loop protecting the RBS [10]. The temperature response of these thermometers was designed on the basis of the melting temperature of the minimum free energy structure, with an increased stem length, a smaller hairpin loop, or a reduction in number of bulges resulting in an increase in the melting temperature [9,10]. More recently, RNA thermometers have also been designed to be heat-repressible using an RNase E-mediated mechanism [4]. These previous studies have provided important results towards an elucidation of mechanisms underlying RNA thermometer operation, modulation of their response to temperature, and towards prediction of their behaviour.

There are at least three striking aspects about the functioning of RNA thermometers. First aspect is the startling complexity in the secondary structure of the naturally occurring RNA thermometers in comparison to the synthetic ones. Second aspect is the multitude of different secondary structures possible, in addition to the minimum free energy structure, that an RNA thermometer can adopt at any given temperature. The activity of a thermometer is directly dependent on the proportion of these different structures and their respective activities. Third aspect is how even a one base-pair change in sequence can qualitatively change the temperature response. In particular, among the published synthetic thermometers [10], a one base-pair change appears to shift the response from linear-like to switch-like (U2 and U9 in [10]). The role of these factors in determining the threshold value and sensitivity of an RNA thermometer is generally unclear (Fig. 1B).

**Figure 1.**
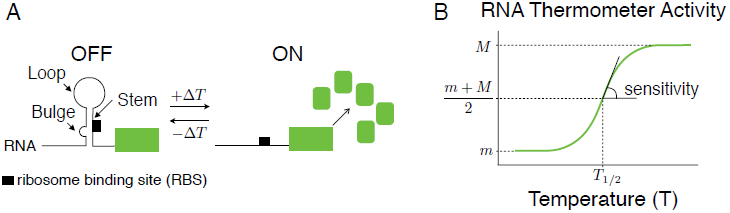
Temperature response of a RNA thermometer. A. Illustration of the functioning of a simple RNA thermometer. Structural features such as loops, bulges, and stems are indicated. B. Green solid line represents the activity of the RNA thermometer as a function of temperature. Key quantitative feature of the response such as threshold and sensitivity are emphasized.

Here, our objective is to design a set of RNA thermometers with different sensitivities in their response to temperature. We used a combination of experimental measurements in cells and in cell-free biomolecular ‘breadboards’ as well as computations of RNA secondary structures to achieve this objective. We studied a set of existing synthetic thermometers, finding consistency with existing results and among our experimental and computational analyses. Next, we designed a library of thermometers by systematically changing a thermometer sequence one base pair at a time, finding, computationally, that a range of temperature responses are possible. Finally, we constructed this library and found a variety of largely-linear responses in the temperature range 29°C–37°C, with different starting values, fold-changes, and slopes. When assessed against the computational expectations, we find that, while the individual measurements do not correlate strongly, the systems-level trends match well. These results should help in developing a toolbox of temperature-regulatory biomolecular circuit components for synthetic biology applications.

## 2 Results

### 2.1 Analysis of existing RNA thermometers

We started our investigation with an analysis of two existing synthetic RNA thermometers X and Y (respectively, U2 and U9 from [10]). These differ in sequence by only one base-pair, yet have a qualitatively different response to temperature (Fig. 2A).

**Figure 2.**
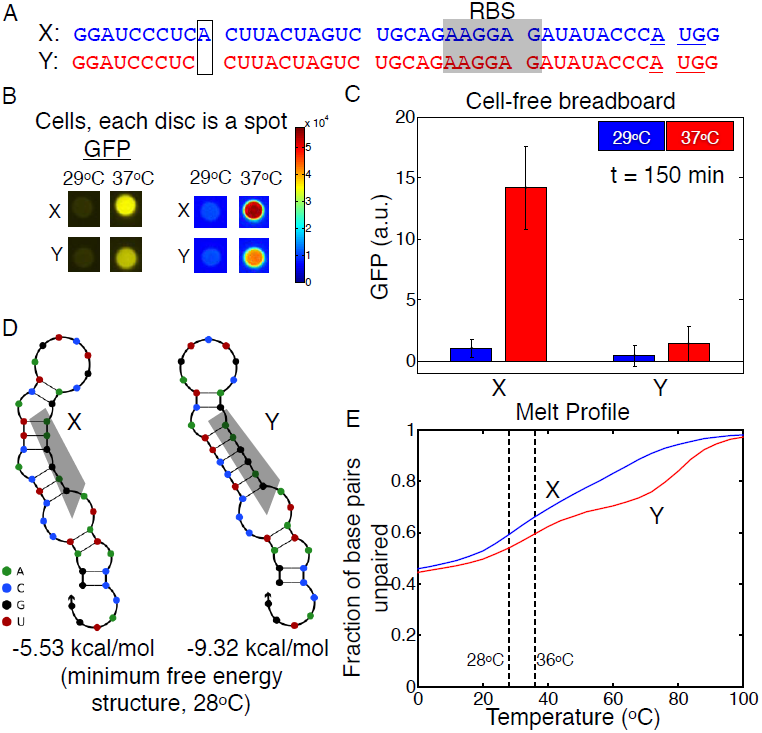
Characterisation of two existing thermometers computationally and experimentally. A. Thermometer sequences. B. Thermometer activity in cell spots. Intensity represents green fluorescence. C. Quantification of thermometer activity in cell-free breadboards. Each bar is a mean of three separate measurements with the error bars representing one standard deviation. D. Secondary structures of the minimum free energy structure and corresponding free energies. E. Melt profiles of the thermometers.

*Measurements in cells:* To get an initial estimate of thermometer activity as a function of temperature, plasmids [10] containing either of these thermometers in the 5’-UTR region of a gene coding for a green fluorescent protein (GFP-trps16) were transformed into *E. coli* (JM109). These cells were then spotted onto multiple LB-agar plates containing 100*μ*g/ml carbenicillin and incubated overnight at different temperatures — 29°C and 37°C. Subsequently, these colonies were imaged under UV light in a fluorescence imager (Fig. 2B). We found that the fluorescence of the colonies increased as a function of temperature, consistent with previous results. Further, there were differences between X and Y in terms of the extent of increase, with cells with the X construct being more fluorescent than the cells with the Y construct at higher temperatures, also consistent with previous results.

*Measurements in a cell-free biomolecular breadboard:* For a more quantitative estimate, we used an *E. coli* cell-extract-based biomolecular breadboard, which allows transcription and translation in a rapid prototyping platform [14]. Plasmid DNA (5nM) was incubated in the breadboard at different temperatures — 29°C and 37°C — and the GFP produced was measured at *t* = 150 minutes. All breadboard measurements were performed in black transparent-bottomed 384-well plates (Perkin Elmer) and a platereader (Synergy BioTek 1), with fluorescence excitation and emission set to 485nm and 515nm, respectively. We found that the thermometer activities increased with temperature (Fig. 2C). Further, the activity of thermometer X increased more than that of Y. These are consistent with expectations based on published results and with the cellular measurements presented above.

In addition to the affect of temperature on the secondary structure of the RNA thermometers, it is likely that temperature affects other aspects of the measurement assays described above. These include the fluorescence parameters of GFP, the activity of RNA polymerase during transcription, and the growth rate of cells. However, these factors should affect both thermometers in a similar fashion. Therefore, in our investigations, we focus on the relative difference in thermometer activity.

*Computations of RNA thermodynamics:* To assess the similarity between these measurements and theoretical expectations based on thermodynamic considerations, we used the computational web server NUPACK [16]. Given an RNA sequence, NUPACK can compute the minimum free energy and the corresponding structure at different temperatures. We computed these structures at 28°C using the sequences of X and Y and found that they were largely similar (Fig. 2D). The major difference is the presence of a bulge in the stem containing the RBS in X. Further, the free energy of the X structure is larger than that of Y. These suggest that the thermometer X should be less stable than Y, and consequentially that expression of GFP controlled by X should be higher than that of GFP controlled by Y at each temperature. This is consistent with results established previously and with the experimental measurements presented above.

For a given temperature, in general, there are multiple structures that an RNA molecule can adopt. These other structures may also contribute to the activity of an RNA thermometer. To take these into account, we used NUPACK to calculate the fraction of unpaired base pairs as a function of temperature. As the thermometer activity is proportional to the extent of its melting, we used this fraction as another measure of its activity. Comparisons of the melt profiles of X and Y show that X melts more than Y at each temperature (Fig. 2E). These are also consistent with our expectations.

Together, these results analysing existing RNA thermometers, both experimentally and computationally, are consistent with published results and with each other.

### 2.2 Design of a library of RNA thermometers

Noting that the two thermometers studied above, with different responses, differed in sequence by only one base pair, we wondered whether other one base pair changes to the thermometer sequence could generate the desired diverse set of temperature responses. There are 43 base pairs in the thermometer X, with a deletion at the 10^th^ position giving the thermometer Y (Fig. 3A). Other one base-pair changes include a deletion at any other location as well as replacing the existing nucleotide with the other three possibilities, a total of 4 × 43 = 172 possibilities. For reasons of scale, we focussed on the 10^th^ position as well as the position involved in the secondary structure base-pairing of the minimum free energy structure. This gives rise to a library of 4 × 19 = 76 variants (Fig. 3A, listed in Supplementary information A).

**Figure 3.**
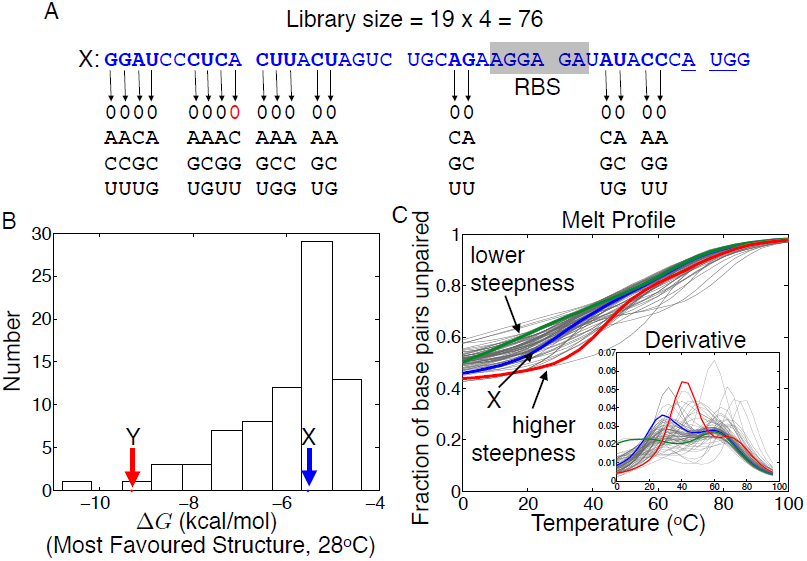
Computational analysis of a thermometer library. A. Sequence profile of the thermometer library. The starting sequence, of thermometer X, is given in blue, with the base pairs to be mutated in boldface. Red base pair indicates the change that gives thermometer Y. Black base pairs represent the base pair changes required to generate the library. B. Histogram representing the free energy of the minimum free energy structure at 28°C. The locations of the thermometers X and Y are highlighted using arrows. C. Melt profiles of thermometer library. Blue line represent the melt profile of thermometer X. Light grey lines represents the melt profiles of remaining thermometers. Red and green lines are highlighted thermometers whose melt profiles show, relative to thermometer X, a lower and higher steepness, respectively. Inset shows the derivatives of the melt profiles computed through a first difference.

*Computational analysis suggests a diversity of responses:* To check if these different thermometers could indeed generate a diversity in responses desired, we used NUPACK to compute their free energies and melt profiles. The free energies computed at 28°C show a diversity of values, across the range from those of X and Y (Fig. 3B). This suggests that different responses may be possible in the library. Similarly, the melt profile computations show that a variety of responses could be generated (Fig. 3C). These include those melt curves that are more linear-like compared to that of X as well as more switch-like. Considering the derivative of the melt curves emphasises the diversity in slopes and their threshold values that may be generated (Fig. 3C, inset). These computations suggest that the design of such a library may achieve its desired objective.

### 2.3 Experimental characterisation of thermometer library

We experimentally constructed the library using standard procedures, as presented below.

*Methods:* Green fluorescent protein in pBSU2 was replaced from gfp-Trp16 to deGFP-T500. deGFP-T500 was amplified using PCR and digested with restriction enzymes NcoI and HindIII. Similarly, pBSU2 was digested with these restriction enzymes and further treated with Antarctic phosphatase. These were then ligated with the Quick Ligase Kit (NEB) and transformed.

The desired thermometer sequences were then constructed using PCR. For each thermometer, four sets of primers were required. The first primer was homologous to a sequence approximately 250 base pair upstream of the promoter of the plasmid pBSU2-deGFP. The second primer was homologous to the X thermometer and contained the desired promoter. These two primers were used to PCR the promoter region with some part of the thermometer. The third primer was designed to substantially overlap with the second primer and contained the desired mutation. The fourth primer was designed to bind to a sequence approximately 250 base pair down stream of the terminator of the plasmid pBSU2-deGFP. These two primers were use to PCR the green fluorescent protein and part of the thermometer region. Finally, these two pieces were fused using the first and the fourth primer. All primers were ordered from IDT Inc. (San Diego, California, USA) and their respective sequences are listed in Supplementary Information B. All PCR reactions were carried out using PhusionΩ Hot Start Flex 2X Master Mix (NEB).

Finally, after PCR, these thermometers were ligated into the vector backbone of pBSU2. For this, these PCR products were mixed, and digested with the restriction enzymes SacI and HindIII. A mixture of all PCR products, when run on a gel, showed the right size of the product (Supplementary Information C). Similarly, the vector pBSU2-deGFP was digested and additionally treated with Antarctic Phosphatase (NEB). This mixture of thermometers and the vector were ligated using the Quick Ligase Kit (NEB) and transformed into chemically competent *E. coli* JM109 cells. The obtained colonies were screened by sequencing using the M13 primers. After one round of screening, we recovered 38 different variants from the library. Additionally, one of the screens was found to be the original thermometer. These 39 thermometers were used for further measurements.

*Measurements:* For a quantitative assessment of temperature responses of these thermometers, we measured the thermometer activities in molecular breadboards, as described above, at three different temperatures — 29°C, 33°C, and 37°C (Fig. 4A, B). We find a variety of responses different from the starting response, even though various responses arise from a thermometer sequence that differs from the starting thermometer sequence by only one base-pair. In particular, while all the responses increase with increasing temperature, the extent of increase relative to temperature was different. To quantify different features of this library better, we computed the maximum fold change and the slopes of the response in the given temperature range (Fig. 4C, D). We find that the maximum fold change is largely similar even though the basal activity, the activity at 29°C, changes (Fig. 4C). Further, there is a slight downward trend in the maximum fold-change as the basal activity increases. Finally, based on the computed slopes, we find that the responses are largely linear in the given temperature range with a variety of slopes (Fig. 4D). These results show that there is a diverse range of thermometer profiles in the library, as required from the aim of constructing this toolbox.

**Figure 4.**
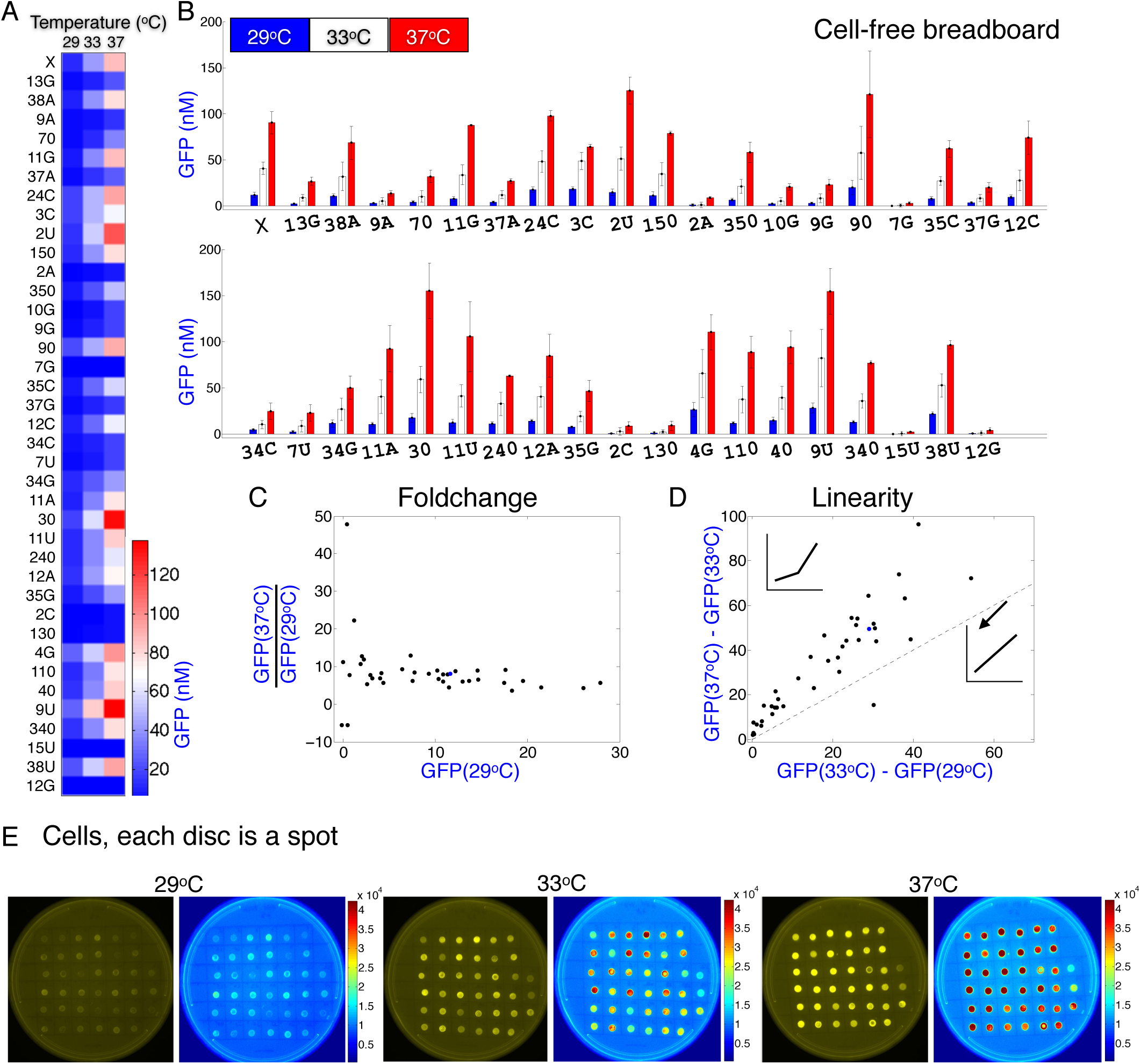
Experimental measurements of the thermometer library show a variety of responses. A. Rows represent activity levels of different thermometers. The activity levels are mean of three separate measurements. These values are normalised using a GFP calibration performed at 29°C. B. Replotting of data from A. Each set of three bars represents the activity levels of different thermometers. Blue, white, and red represent the temperatures 29°C, 33°C, and 37°C, respectively. The height of the bars corresponds the mean levels of three separate measurements with the standard deviation shown as the error bar. C & D. Each black dot represents a separate thermometer. A blue dot is used for the starting thermometer X. E. Thermometer activity in cell spots. Intensity represents green fluorescence. The identity of spots, read in a left-right, top-bottom manner, is the same as in A.

To get an estimate of the temperature responses of these thermometers inside cells, we spotted them on LB-agar plates and grew them overnight at 29°C, 33°C, and 37°C. These plates were then imaged with a fluorescence imager (Fig. 4E). The fluorescence values in these measurements, in general, increased with temperature. There were differences, however, in the rate of increase. For example, consideration of the fluorescence values at 33°C highlights the differences in responses that can be obtained. Therefore, this thermometer library can function inside cells and display a diversity of responses to temperature.

### 2.4 Assessment of match between computations and experiments

We have experimentally measured the thermometer responses at three different temperatures and interpreted these measurements in terms of the key response characteristics. From a computational standpoint, we had used two quantitative characteristics — free energies and melt profiles — to check the diversity in the library. Finally, we assessed the match between these computational and experimental results.

For the comparison with the free energy of the minimum free energy structure, we plotted the free energy against experimentally measured expression level for each thermometer and at the different temperatures (Fig. 5A). For a good correlation, we would expect a linear trend, which is not seen. At best, the cluster of points may be seen to appear to be below a straight line, in a triangular-shaped region. In contrast to the weak correlation observed here, the overall trend of the fold change plot (Fig. 5B) is of decreasing nature, similar to that obtained experimentally (Fig. 4C). To obtain the computational version of the fold change plot, we computed the difference in the free energies of each thermometer and plotted it against the free energy at the lowest temperature. Similarly, the slope plot corresponding to the free energies, where the difference in free energies at the successive temperatures are plotted against each other (Fig. 5C), bears the signatures of a linear response, correlating well with the experimental one (Fig. 4D). Therefore, the overall trends appear to match well, but the individual values do not.

**Figure 5.**
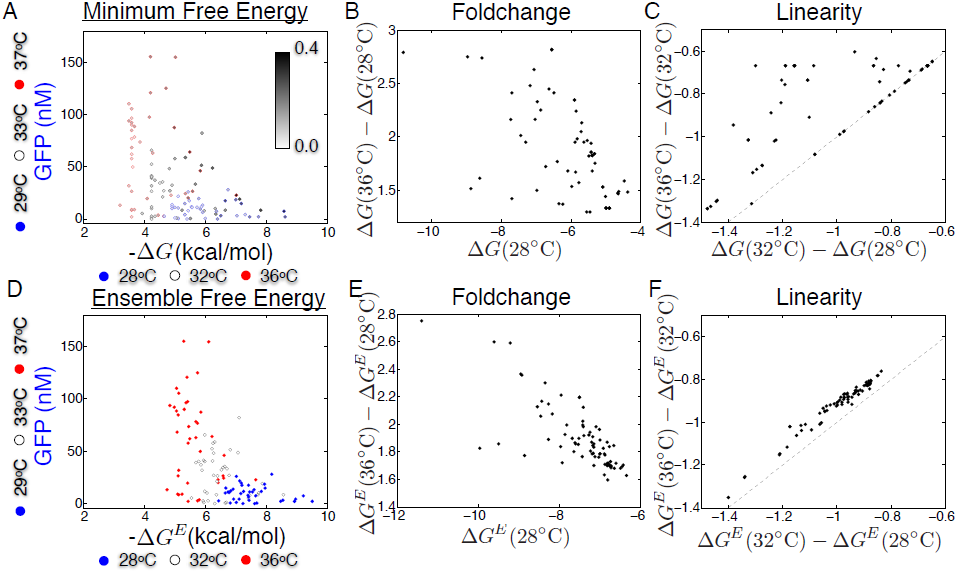
Assessment of experimental measurements and computations. A. Each dot represents an individual thermometer plotted with its experimentally measured value on the Y-axis and the free energy of the minimum free energy structure on the X-axis. Blue, white and red dots represent, respectively, the temperatures 29°C, 33°C, and 37°C for experimental measurements and the temperatures 28°C, 32°C, and 36°C for the computations. Each dot is shaded according to the equilibrium probability of the minimum free energy structure. B. Each dot represents an individual thermometer plotted with its net difference in free energy of the minimum free energy structure between the temperatures 28°C and 36°C on the Y-axis and the free energy of the minimum free energy structure at 28°C on the X-axis. C. Each dot represents an individual thermometer with its net difference in free energy of the minimum free energy structure between the temperatures 32°C and 36°C on the Y-axis and the net difference in free energy of the minimum free energy structure between the temperatures 28°C and 32°C on the X-axis. Dashed line represents where the dots would lie if the response was linear in this temperature range. D. Each dot represents an individual thermometer plotted with its experimental value on the Y-axis and the free energy of the ensemble on the X-axis. Blue, white and red dots represent, respectively, the temperatures 29°C, 33°C, and 37°C for experimental measurements and the temperatures 28°C, 32°C, and 36°C for the computations. E. Each dot represents an individual thermometer plotted with its net difference in ensemble free energy between the temperatures 28°C and 36°C on the Y-axis and tine ensemble free energy at 28°C on the X-axis. F. Each dot represents and individual thermometer with its net difference in ensemble free energy between the temperatures 32°C and 36°C on the Y-axis and the net difference in ensemble free energy between the temperatures 28°C and 32°C on the on the X-axis. Dashed line represents where the dots would lie if the response was linear in this temperature range.

A possible reason underlying the discrepancies observed is that the free energy considered is that of the minimum free energy structure, which may be a small fraction of the overall ensemble of structures. To address this, we shaded the dots in Fig. 5A according to the fraction of the minimum free energy structure. These computations show that the minimum free energy structure may be only a small fraction of the overall ensemble. To investigate this further, we considered the free energy of the entire ensemble and assessed them against the experimental measurements. We find similar trends (Fig. 5D–F). The individual thermometer responses do not show strong correlations, while the overall trends match.

For comparison with the melt profile, we first plotted the experimental measurements against the melt profile values at each temperature (Fig. 6A). The plot does not reveal a strong correlation. On the other hand, when the fold-change and linearity of the response is assessed from the melt profiles, there is a striking correlation with the observed trends (Fig. 6B–C). These plots were with the melt profile of the entire sequence. To check if a different part of the sequence gives a better match, we considered the average equilibrium probability that the base pairs belonging to the RBS are unpaired. For this we computed the average probability, and analysed it, both in terms of the correlation with the experimental data (Fig. 5D) as well as for the trends within it (Fig. 6E–F). As above, we find no strong match with the individual experimental datapoints, although the overall trends, especially of the slope response, match reasonably well.

**Figure 6.**
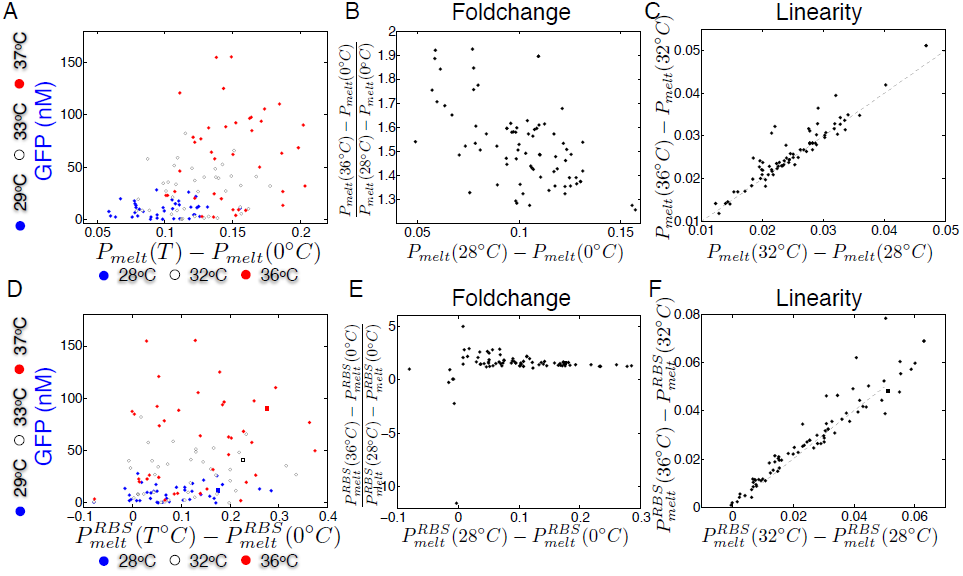
Assessment of experimental measurements and melt profile computations. A. Each dot represents an individual thermometer plotted with its experimental value on the Y-axis and the extent of melt on the X-axis. Blue, white and red dots represent, respectively, the temperatures 29°C, 33°C, and 37°C for experimental measurements and the temperatures 28°C, 32°C, and 36°C for the computations. B. Each dot represents an individual thermometer plotted with its net difference in melt profile between the temperatures 28°C and 36°C on the Y-axis and the extent of melt at 28°C on the X-axis. C. Each dot represents and individual thermometer with its net difference in melt profile between the temperatures 32°C and 36°C on the Y-axis and the net difference in melt profile between the temperatures 28°C and 32°C on the on the X-axis. Dashed line represents where the dots would lie if the response was linear in this temperature range. D. Each dot represents an individual thermometer plotted with its experimental value on the Y-axis and the extent of melt of the RBS region on the X-axis. Blue, white and red dots represent, respectively, the temperatures 29°C, 33°C, and 37°C for experimental measurements and the temperatures 28°C, 32°C, and 36°C for the computations. E. Each dot represents an individual thermometer plotted with its net difference in melt profile of the RBS region between the temperatures 28°C and 36°C on the Y-axis and the extent of melt of the RBS region at 28°C on the X-axis. F. Each dot represents and individual thermometer with its net difference in extent of melt of the RBS region between the temperatures 32°C and 36°C on the Y-axis and the extent of melt of the RBS region between the temperatures 28°C and 32°C on the on the X-axis. Dashed line represents where the dots would lie if the response was linear in this temperature range.

Overall, through all the metrics considered, we note that the individual values themselves do not exhibit a strong correlation, but the characteristic trends are similar. Therefore, while these computational considerations were useful in designing the library, the exact correspondence between the computations and experiments is not present, perhaps because of the assumptions in the computations such as the choice of parameters and restricted set of possible structures considered.

## 3 Discussion

Temperature-sensing RNA thermometers can have multiple applications. Using a combination of experimental measurements, in cell-free biomolecular breadboards and in cells, and computations of RNA thermodynamics such as free energies, minimum free energy structure, and melt profiles, we have presented the following three results. First, we found broad agreement with our methodologies and previous measurements on a set of thermometers. Second, we computationally found that a wide array of temperature responses are possible in a library of thermometers, each member of which is a one base pair alteration of a starting thermometer sequence. Third, we synthesized such a library and found a diversity of responses with different sensitivities. The overall trends of the responses obtained computationally and measured experimentally match reasonably well, although the individual values do not show a strong correlation.

It is interesting that even a one base pair change in the sequence of an RNA thermometer can generate a substantial change in the temperature response. We have observed this in our results both computationally and through the experimental measurements. It is likely that relatively complicated secondary structures that are found in the natural contexts may have evolved so that sensitivity to single base pair alterations is minimised.

An immediate task for future work is to correlate the experimental measurements in the cell-free breadboards with quantitative measurement in cells. As a next step, it may be useful to demonstrate functional applications of these thermometers, particularly in the development of circuits that need to interface with temperature. An example of this is to use these as the input stage of a pulse generating incoherent feedforward loop circuit, thereby generating a pulse of gene expression triggered by temperature.

The design and development of new biomolecular circuits and components, often based on naturally occurring ones, facilitates various synthetic biology applications, such as in metabolic engineering, agriculture, and medicine [15]. RNA is an attractive substrate for this, due to its substantial regulatory and sensing potential [2,3] as well as the availability of computational tools to estimate its behaviour [1,13]. Indeed, RNA molecules can be designed to sense extremely specific signals such as metabolites or other RNAs [7] as well as more global signals such as temperature [8]. Here, we have presented experimental and computational results for the development of RNA-based temperature sensors. In addition to the multiple applications mentioned above, such as in large bioreactors, for thermal imaging and control of gene expression, as well as for designing temperature robustness, these results highlight the usefulness of RNA for synthetic biology.

## Acknowledgements

We thank Prof. Dr. Ralph Bock and Dr. Juliane Neupert from Max-Planck-Institut fur Molekulare Pflanzenphysiologie, Potsdam, Germany, for their kind gift of plasmids, including pBSU2 and pBSU9. Finally, we thank Dr. Harry Choi, Clare Hayes, Anu Thubagere, and Dr. Jongmin Kim for being gracious with their time and guidance. This work was supported in part by the Defense Advanced Research Projects Agency (DARPA/MTO) Living Foundries program, contract number HR0011-12-C-0065 and the National Science Foundation award number 1317694.

